# Batch Equalization with a Generative Adversarial Network

**DOI:** 10.1101/2020.02.07.939215

**Authors:** Wesley Wei Qian, Cassandra Xia, Subhashini Venugopalan, Arunachalam Narayanaswamy, Jian Peng, D. Michael Ando

## Abstract

Advances in automation and imaging have made it possible to capture large image datasets for experiments that span multiple weeks with multiple experimental batches of data. However, accurate biological comparisons across the batches is challenged by the batch-to-batch variation due to uncontrollable experimental noise (e.g., different stain intensity or illumination conditions). To mediate the batch variation (i.e. the batch effect), we developed a batch equalization method that can transfer images from one batch to another while preserving the biological phenotype. The equalization method is trained as a generative adversarial network (GAN), using the StarGAN architecture that has shown considerable ability in doing style transfer for consumer images. After incorporating an additional objective that disentangles batch effect from biological features using an existing GAN framework, we show that the equalized images have less batch information as determined by a batch-prediction task and perform better in a biologically relevant task (e.g., Mechanism of Action prediction).

## 1 Introduction

A key challenge in identifying relationships within the high-resolution images of high-content biological assays is minimizing the influence of technical sources of variation arising from sample handling. Extensive experiments, especially with biological replicates, are often run over multiple weeks, with each week’s smaller experiment forming one batch of data. Changes in both the initial state of the cell population as well as the inevitable deviations during assay preparation and imaging can lead to substantial differences in the appearance of the cells in images. This is termed batch effect (Leek et al., 2010; Caicedo et al., 2017). While existing computer vision models are helping us understand our data better, they are still easily deceived by the batch effect. The batch to batch variation can overpower the actual biological variation (Figure 1a), especially in weak/un-supervised settings (Pawlowski et al., 2016; Ando et al., 2017). In their analysis of various datasets, Ando et al. (2017) and Venugopalan et al. (2019) show that much of the variance in the data is, in fact, capturing the batch information instead of the biological information. Since it is often not possible to run the ideal experiment with all the experimental conditions in one batch, it is crucial to address the batch effects for our computation models to capture the right biological features.

**Figure 1:**
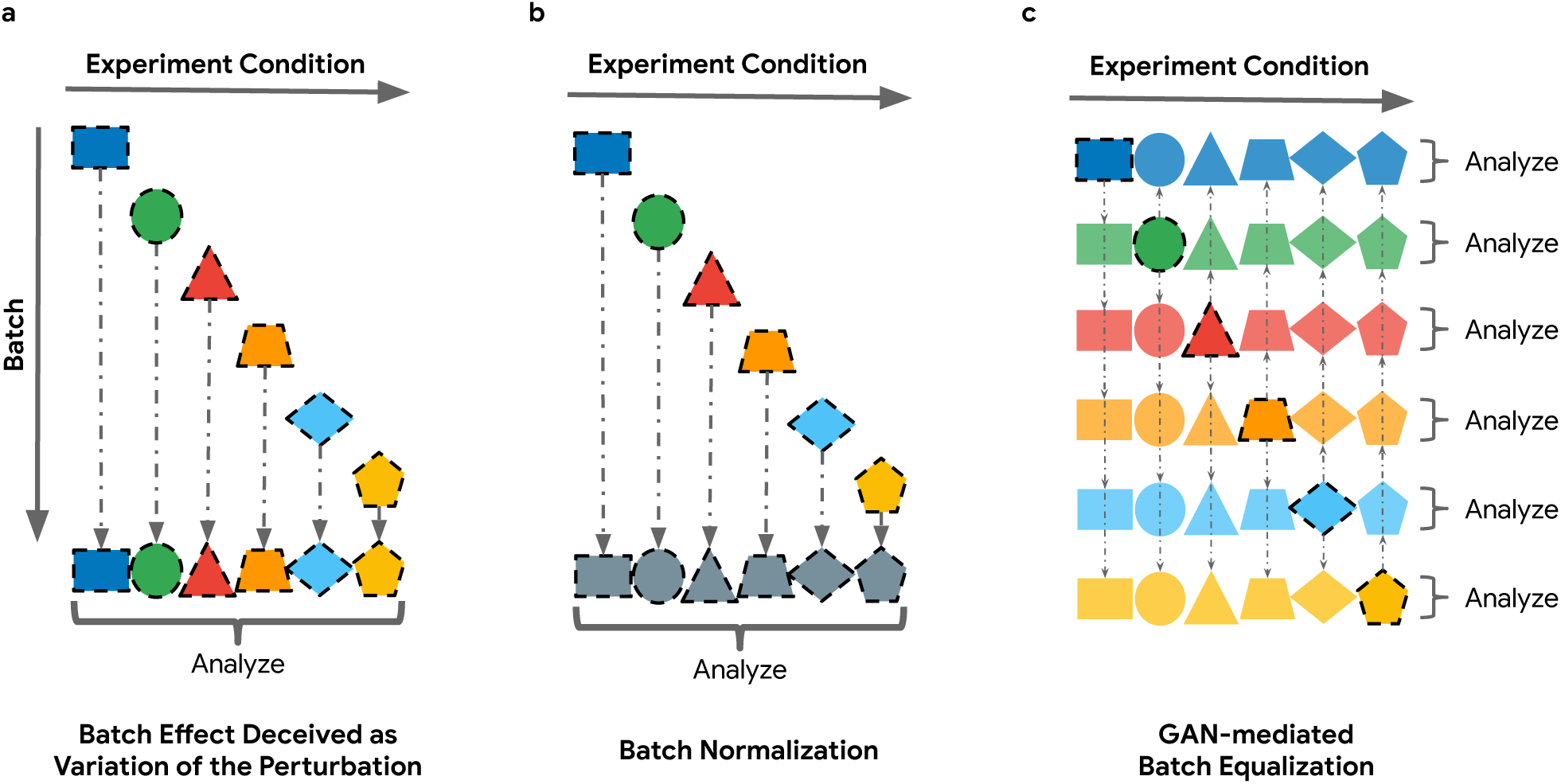
Normalizing batch effect through equalization. Different colors represent the variation caused by the batch effect, and different shapes represent different biological features as a result of the experiment condition difference. a) Batch effect can deceive the computation model where the batch variation could overpower the biological variation that we want to understand. b) Existing method tries to mediate the batch effect through normalization. c) In this work, we propose a generative model that learns to transfer images from the distribution of one batch to another without modifying the biological phenotype giving us an ideal dataset where all experiment conditions are run in all batches and mediating the batch effect naturally.

Even though laboratory automation, along with iterating and optimizing the experiment protocol, can mediate batch effect, there is a ceiling for improvement due to the intrinsic biological variability and technical execution. Therefore, many analysis and computation model include a batch normalization step (Figure 1b). In Pawlowski et al. (2016), heuristics like illumination correction and grey-scaling are applied to the images to mediate the batch effect. In the down-stream analysis, high dimension image data is often projected into lower dimension space through heuristic algorithms or neural networks as embedding. Therefore, many works have been developed to normalize the batch effect for the embedding representations. In Kothari et al. (2011), simple statistical normalization is used to correct the batch effect for the embedding vectors, and these include mean, rank, and ratio normalization as well as parametric, or non-parametric Bayesian prior. In Ando et al. (2017), a statistical tool Typical Variational Normalization (TVN) is introduced to re-scale the variation for the embedding based on the variation analysis from the negative control. In Tabak et al. (2017), a domain adaptation framework is introduced where the domain-specific information is removed from the embedding representations by minimizing a loss function based on Wasserstein distance. Unlike Tabak et al. (2017) where domain-specific information is ‘removed’, Amodio et al. (2018) learn a transformation (or ‘neural edit’) in the latent space of an embedding vector autoencoder such that embedding in one domain can be transferred to an embedding in another by adjusting the latent space of the autoencoder.

While promising results have been achieved with these embedding normalization techniques, we do not have much understanding of what they are doing in embedding space. These normalization methods are potentially altering the right biological features (e.g., cell size/density), and we hope to bring greater interpretability and intuition to the normalization process by mediating the batch effect in the image pixel space. In addition, image-level batch effect mediation can be easily combined with embedding level normalization such that we can take advantage of the benefit from both worlds. While existing image preprocessing pipelines often include some simple heuristic for image normalization, they are often too simple to handle the artifacts from the batch effect. For instance, it’s common to compute a median (or other percentiles) image as a proxy for the “flatfield” image. However, such a heuristically-chosen “flatfield” image might not be the right image for all images, and therefore, we are interested in a learning-based method that can make the “right” image on a real per-image basis.

Recent advances in Generative Adversarial Network (Goodfellow et al., 2014) have demonstrated astonishing abilities at mapping images from one “style” to another (i.e., style transfer). For instance, Pix2Pix (Isola et al., 2017) leverages Conditional GAN (Mirza and Osindero, 2014) and learns to transfer image style pixel by pixel in a supervised setting with paired images. However, obtaining data pairs for Pix2Pix training is expensive and sometimes impossible (the same experiment cannot be done in two different time points). Therefore, CycleGAN (Zhu et al., 2017) and DiscoGAN (Kim et al., 2017) were later developed to circumvent the need for paired data. Both architectures learn to preserve semantic information while transferring style from one image to another. A cycle consistency loss requires the model to retain the semantic information of the source image through a cycle of transformation such that the transformed image can be converted back to its original form.

If we treat each batch as an individual style, we can formulate the batch normalization problem as a batch equalization problem where we learn to transfer all images to the same batch with the biological (or semantic) feature intact such that the batch effect can be mediated naturally (Figure 1c). In this work, we leverage StarGAN (Choi et al., 2017), a multi-domain/style extension of CycleGAN, and train a generator to transfer negative control from one batch to the style of another batch. To further disentangle the biological features (e.g., cell size/density) from the image features (e.g., illumination), we introduce additional regression tasks to make sure the biological evidence is intact during the transformation. By equalizing the images with our batch equalization GAN, we can reduce the batch variation in the images while the biological features still correlated with the treatment conditions remained intact.

## 2 Methods

### 2.1 Overall framework

To transfer image **X** from its original batch to another batch *t*, we want to learn a conditional generator *G* : (**X**, *t*) ↦ **X**′ where **X**′ looks like a real image from batch *t* while keeping the original biological features as **X**. Since we do not have training labels for **X**′ either, our generator *G* is trained with a discriminator model *D* serving as the teacher for *G*. Parameterizing both *G* and *D* as neural networks with *θ*_*G*_ and *θ*_*D*_, we can train them together in the GAN setting, as shown in Figure 2. First of all, given an image, the discriminator will predict if the image is a generated image or a real image (*D*_src_). At the same time, the generator will try to fool the discriminator as much as possible by generating a realistic-looking image. In addition, the discriminator will also learn to predict the batch of a given image (*D*_batch_), and the generator is tasked to generate an image *G*(**X**, *t*) that the discriminator will classify as batch *t* such that the generated image will match the real image distribution for batch *t*. To avoid trivial solutions and model collapse, a reconstruction task is introduced to make sure the information content of the image remains intact as much as possible. After the generator transfer image from batch *t*_*i*_ to *t*_*j*_, the generated image also needs to be transferred back to its original batch *t*_*i*_ such that ‖*G*(*G*(**X**, *t*_*j*_), *t*_*i*_) − **X**‖_1_ is minimized. Finally, to disentangle potential biological features that are correlated with the batch variation, the discriminator also learn to predict the entangling biological features such as cell foreground fraction (*D*_entangled_), and the generator learn to disentangle the biological features from the rest of the representation by generating an image with the original biological features determined by the discriminator. After training the generator in the GAN framework with the discriminator, we can transfer the entire image dataset to the same batch target *t* using the generator model *G* such that the batch effect is equalized/mediated and downstream analysis can then be performed on this *generated* ideal dataset.

**Figure 2:**
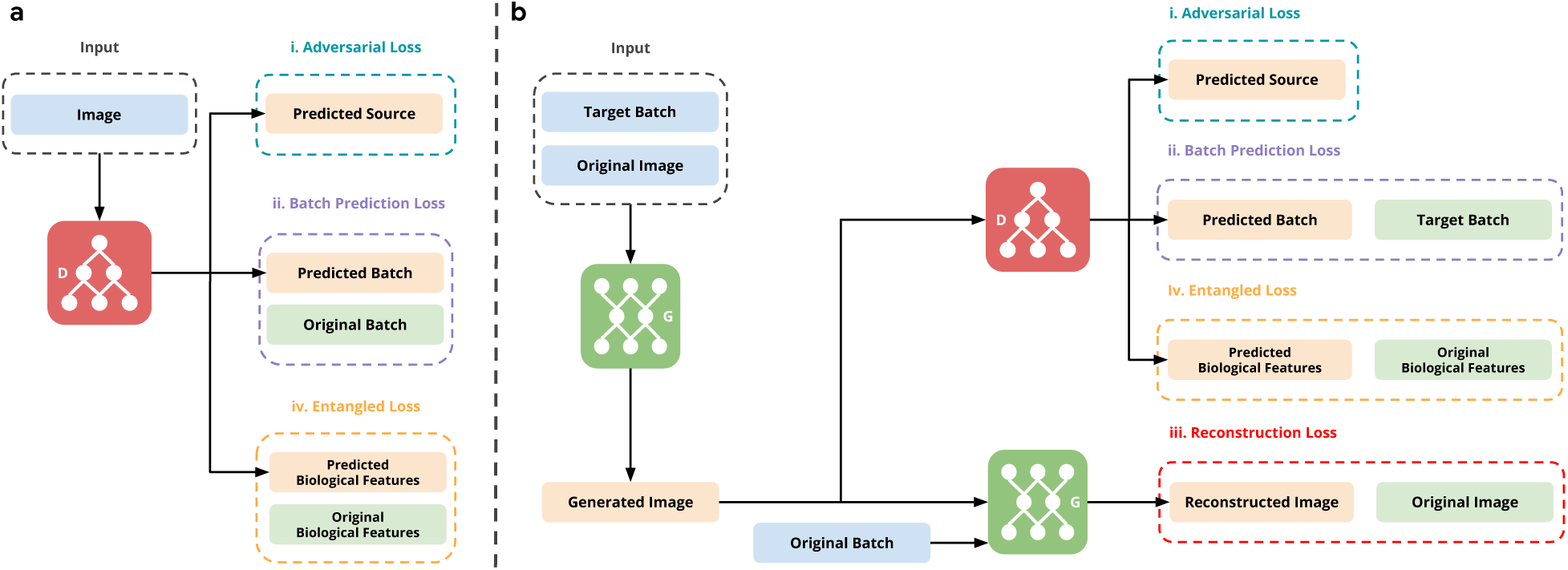
Training the generator and the discriminator in a StarGAN setting (Choi et al., 2017). a) Training procedure for the discriminator *D*. The discriminator *D* takes an image (either real or generated) and predicts its source (real/generated), which batch it comes from, and biological features such as cell size. The adversarial loss helps it to separate real images from generated images. Through the batch prediction loss, it learns to predict the batch of a real image accurately. Finally, we introduce an entangled loss where the discriminator *D* learns to predict the biological features of an image. b) Training procedure for the generator *G*. The generator *G* takes an image and a target batch as input and produces a generated image belong to the target batch. Besides the standard adversarial loss where the generator *G* learns to generate realistic-looking images, *G* also learns to change the batch effect of an image through the batch prediction loss. To avoid model collapse, the original StarGAN framework also has a reconstruction loss encouraging the generated images to keep enough information for the original images - although the apparent semantic content may change. Finally, our new entangled loss is imposed to make sure the biological features for both the generated images and the original images match, so that the semantic content and the biological features remain intact.

### 2.2 StarGAN for batch transformation

Since our framework heavily relies on the original StarGAN (Choi et al., 2017), we will first do a brief review of the StarGAN formulation, and introduce our extension for representation disentanglement in the following section. Under the GAN framework, our discriminator *D* will learn to predict whether an image is a real image from the original data distribution or a generated image from *G*. This is the image source prediction *D*_src_(**X**). While the discriminator is trained to predict *D*_src_ as accurately as possible, the generator is trained to generate images that match the original data distribution as much as possible such that the discriminator cannot accurately predict *D*_src_. This forms the adversarial loss ℒ_adv_, and as both *G* and *D* improve during the training, we can generate image looks more and more realistic. Specifically, we use an improved version of the Wasserstein-GAN (Gulrajani et al., 2017) as the adversarial loss:

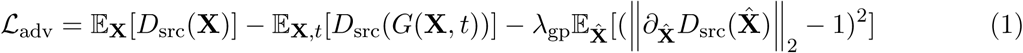

where 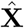 is an interpolation of a real image **X** and the corresponding generated image *G*(**X**, *t*), *t* is sampled from a uniformed distribution over all the target batch, and *λ*_gp_ is the gradient penalty coefficient. The gradient penalty term is introduced to enforce a Lipschitz constraint such that we have a smoother gradient to stabilize the training. Since *G* and *D* are working against each other in this setting, *G* is trained to minimize ℒ_adv_ while at the same time *D* is trained to maximize ℒ_adv_.

In order to make the generated image match the image data distribution of the target batch *t*, the discriminator is tasked to accurately predict the batch of the real images as a distribution overall all the target batch *D*_batch_(**X**). This brings us to a simple cross-entropy loss for the discriminator:

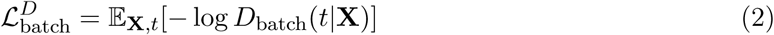

where **X** is sampled from all the images in batch *t*. As the discriminator is trained to predict the batch of an image, the generator’s goal is to transfer an image **X** to batch *t* such that the independently trained discriminator also think the transferred image is from the target batch (i.e., *D*_batch_(*t*|*G*(**X**, *t*)) = 1). Therefore, we can set up a similar cross-entropy loss but for the images from the generator:

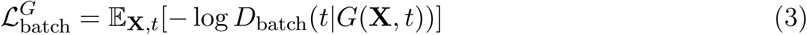

where **X** is sampled from the entire dataset, and *t* is sampled uniformly from the target batch set.

While training *G* to optimize ℒ_adv_ and ℒ_batch_ can give us realistic images look like the ones in the target batch, it is easy to see that the generator can come up with a trivial solution and disregard the content of the original images. Therefore, to force the generator to keep the information content while changing the style of an image, an reconstruction loss is imposed such that after the transformation from batch *t*_*i*_ to *t*_*j*_, the generated imaged, *G*(**X**, *t*_*j*_), must contain enough original information to be transformed back to the original image. The differences between the reconstructed image, *G*(*G*(**X**, *t*_*j*_), *t*_*i*_), and the original image, **X**, are penalized by a *L*-1 norm as the following:

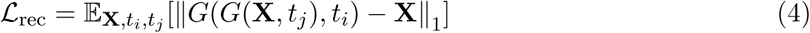

where *t*_*i*_ and *t*_*j*_ are independently and uniformly sampled from the target batch set, and *L*-1 norm is chosen because it tends to generate sharper image than *L*-2 norm as shown in Isola et al. (2017).

### 2.3 Representation disentanglement through regression tasks

While the reconstruction loss ℒ_rec_ in StarGAN aims to keep the semantic content intact, the training still suffers from the feature entanglement problem as discussed in Liu et al. (2018). Among the negative control images, we still observe a significant difference in cell density/size across different batches. Since these biological features are entangled with the batch effect, the generator will still shift these biological features to trick the discriminator resulting in unintended biological modification.

Let **Y**_**X**_ be the entangling biological features we want to remain intact during the batch transformation for image **X**. We want the generator to produce image *G*(**X**, *t*) such that **Y**_**X**_ = **Y***G*(**X**,*t*). **Y** can be given labels such as cell type, or pre-computed label such as foreground fraction of each type of stain. In our case, we use the foreground fraction of DAPI as a proxy for the cell density and the foreground fractions of other stains as a proxy for the cell size.

During the GAN training, the discriminator tries to predict **Y** as accurately as possible for a real image. The discriminator’s predictions are represented as *D*_entangled_(**X**). Since our **Y** are the foreground fractions of each type of stain, we have a regression task and employ a *L*-2 loss:

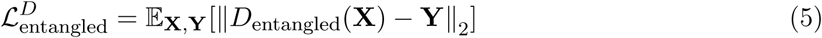

and the corresponding loss for the generator becomes:

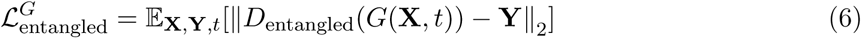

where *t* is sampled uniformly from the target batch set.

Putting everything together, our final objectives for the generator and the discriminator are:

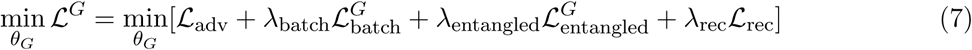

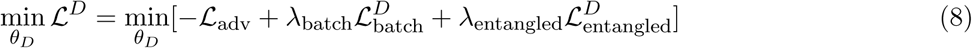

where *θ*_*G*_ and *θ*_*D*_ can be optimized through stochastic gradient descent methods like Adam (Kingma and Ba, 2014). Our implementation details, hyper-parameter selections, and neural network parameterizations for *G* and *D* are described in the supplement section.

### 2.4 Mediating the batch effect

Once the generator *G* is trained, we can transfer all the images in our dataset to the same target batch *t* by *G*(**X**, *t*) (including **X** in batch *t*). Since all the generated images *G*(**X**, *t*) will have a similar batch effect, the batch effect is effectively mediated for downstream analysis. In addition, since there is no requirement to choose a specific target batch *t*, we can randomly select a target batch *t*. Assuming we have a total number of *T* target batches, we can alternatively generate *T* such an ideal dataset and run the downstream analysis for these *T* datasets through averaging or consensus ensembling.

## 3 Data

For all of our studies, we used the BBBC021v1 dataset (Caie et al., 2010), available from the Broad Bioimage Benchmark Collection (Ljosa et al., 2013). The dataset consists of scientific images from ten weeks (i.e., ten batches) of experiments. Each scientific image is composed of three images that represent DAPI, Tubulin, and Actin phenotypes of the MCF-7 wild-type P53 breast cancer cell line. Each week a distinct set of compounds was tested at eight concentrations, using between three and six 96-well plates for the week. From the 38 compounds, there are 103 total treatments in the dataset. Along with the images, the dataset also includes the mechanism of action (MoA) label for each compound, and we are expected to predict the MoA of a compound given the cell responses shown in the images. Each compound is labeled with one of the twelve MoAs.

In addition to the experimental compounds, each plate contains six wells of negative control (DMSO) and positive control (Taxol), making them the only conditions present for every plate and every week. Since positive control only represents a single type of phenotypic response, we avoided training on it since it could bias our image correction toward a particular drug response. Instead, we only trained on the negative controls since all compounds have a phenotype similar to the negative control at very low concentrations, and our results should generalize better as it is always possible to collect the negative control (i.e., unperturbed) phenotype.

For our training, we concatenate the three corresponding stain images (DAPI, Tubulin, and Actin) into a single image with three channels. Along with the images, we also use the batch information for learning batch transformation, and the foreground fractions of the three images for ensuring the biological features remain intact. The batch information is available in the metadata, and the foreground fraction can be easily computed using the Otsu’s method (Otsu, 1979). After the equalization, we used the MoA label prediction to evaluate if the biological features are intact during image transformation.

### 3.1 Image preprocessing and normalization

We performed preprocessing to convert the three 16-bit images into three 8-bit images similarly to Ando et al. (2017), and image level normalization, such as flat-field correction (Singh et al., 2014) is performed (see supplement for details). We want to point out that the image corrections, along with the preprocessing, are intended to correct for batch effect in the existing pipeline. Thus, we believe that any improvement in reducing the batch variations (as shown later) demonstrates the limitations of the standard approaches and the broad applicability of our approach.

### 3.2 Image embedding

In high throughput cellular image analysis, high dimensional image data is often projected to a low dimensional feature space through pre-trained neural networks for downstream analysis (Pawlowski et al., 2016; Ando et al., 2017; Caicedo et al., 2018). For our study, we generated a 64-dimensional embedding from an input RGB image using a Deep Metric Network (Wang et al., 2014) pre-trained on consumer image datasets. Additional details can be found in the supplement section. We note that previous work such as Pawlowski et al. (2016) and Ando et al. (2017) used a per-channel grayscale image as input to the model and then created a single representation by concatenating the per-channel embeddings. However, Caicedo et al. (2018) demonstrates that high-performance can still be achieved with an RGB image, and they additionally have the benefit of being able to detect important biological features like co-localization, which would be impossible from separate representations of the images.

## 4 Results

### 4.1 Approaches compared

**Original** is the first baseline where original images are used for analysis. Since image-level normalization such as flat-field correction (Singh et al., 2014) is already performed for the original images, performance gains from our equalization approach would show the limitation of these heuristic image normalization methods as well as the necessity of a more sophisticated method.

**TVN** is a normalization method to mediate the batch effect for image embedding (Ando et al., 2017). For this baseline, we apply the image embedding methods directly on the original image and run TVN to normalize the image embedding. Since an embedding level normalization does not concern any normalization on the image level, we also show the combined effect of our image equalization method and the embedding level normalization in a later study. Batch * is the approach presented in this paper and represents

**Batch \*** is the approach presented in this paper and represents an equalized image dataset where all images are equalized to batch * using our approach before we run any of the downstream analyses. Because we can equalize the images to any batch, we equalize the entire dataset to the ten different batches and run analysis separately. Since all the images are equalized to the same batch, we hypothesize that the batch effect would be mediated among these equalized images.

### 4.2 Entangled loss extension prevents biological changes

To qualitatively demonstrate the ability of batch equalization GAN, we transfer the same image to all the target batches and show the modification of our generator made in Figure 3. More examples can be found in Figure S1 and Figure S2.

**Figure 3:**
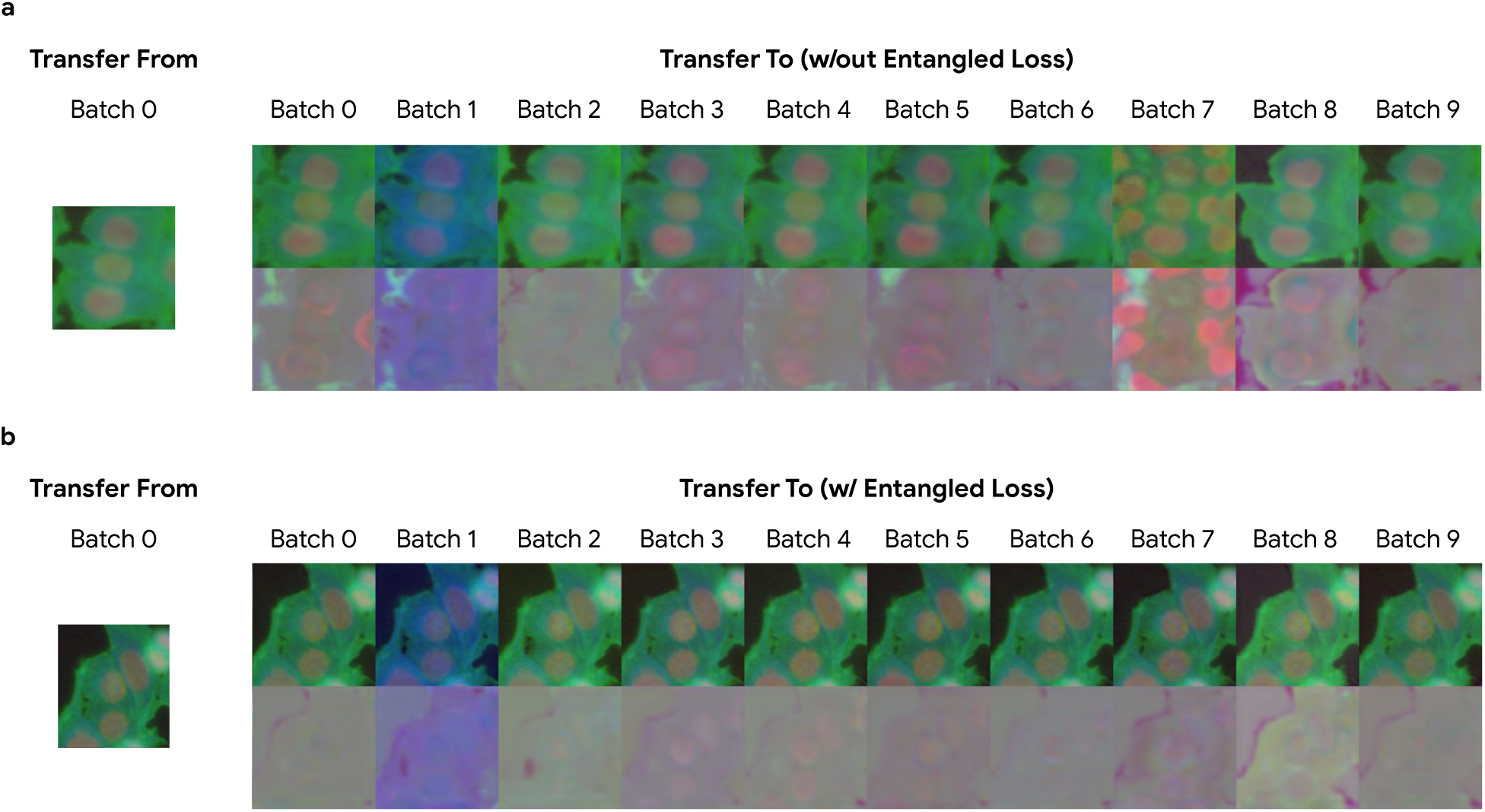
Generator trained in our GAN framework can produce high-quality batch transformation. The first row is the generated image, and the second row is the difference between the generated image and the original image. a) Generator training without the entangled loss ℒ_entangled_ makes dramatic changes to the image with “hallucinated” in Batch 7 while the cell area is decreased in Batch 8. b) After training with the entangled loss, *G* is only transferring the general “style” of the image and keep the content mostly intact.

Training under the original StarGAN framework without the entangled loss ℒ_entangled_, the generator can produce dramatic changes in the image, as shown in Figure 3a. Additional cells are “hallucinated” in Batch 7 while the cell area is decreased in Batch 8. Even though these changes do accurately reflect the batch variation in BBBC021 dataset since the negative controls in batch 7 do have higher confluency than the rest of the batches and the negative controls in batch 8 do have smaller cells. Nevertheless, we don’t want to include them during the batch transformation because these biological features could have true biological consequences. By introducing the entangled loss, we can dramatically reduce these arbitrary changes and keep most of the biological features intact while the model is still able to do sophisticated (and non-linear) updates to model the batch effect, as shown in Figure 3b. The comparison here highlights the necessity of the entangled loss for maintaining the biological features that might be shifted with the target batch in the original StarGAN framework.

We also want to point out that the development of the new entangled loss extension is inspired by visualizing the dramatic effect of the original StarGAN. This again highlights the importance of understanding what has been changed during the normalization stage and why normalizing in the embedding space might not be desirable.

### 4.3 Batch equalization GAN reduces the batch variation in image embedding

To evaluate how much batch information is presented in the image embedding, we used a random forest classifier to predict the original batch of all negative controls from the image embedding. Because of the batch variation in the original image, the classifier can predict the batch of an embedding considerably well. If our batch equalization were successful, we would expect a classifier trained/tested on the equalized dataset to perform worse.

In a five-fold cross-validation study, we compared the batch prediction accuracy of the original image embedding as well as the TVN normalized image embedding to the equalized image embedding. For the equalized image embedding, we transformed the image to each of the target batches and evaluated them individually. As shown in Figure 4a, we see an average of 5% drops in the prediction accuracy after the equalization, even though heuristic-based methods have already been applied to original images to mediate the batch effect. Since TVN is not trained to remove the batch information, we observe more batch predictive power in the TVN normalized image embedding.

**Figure 4:**
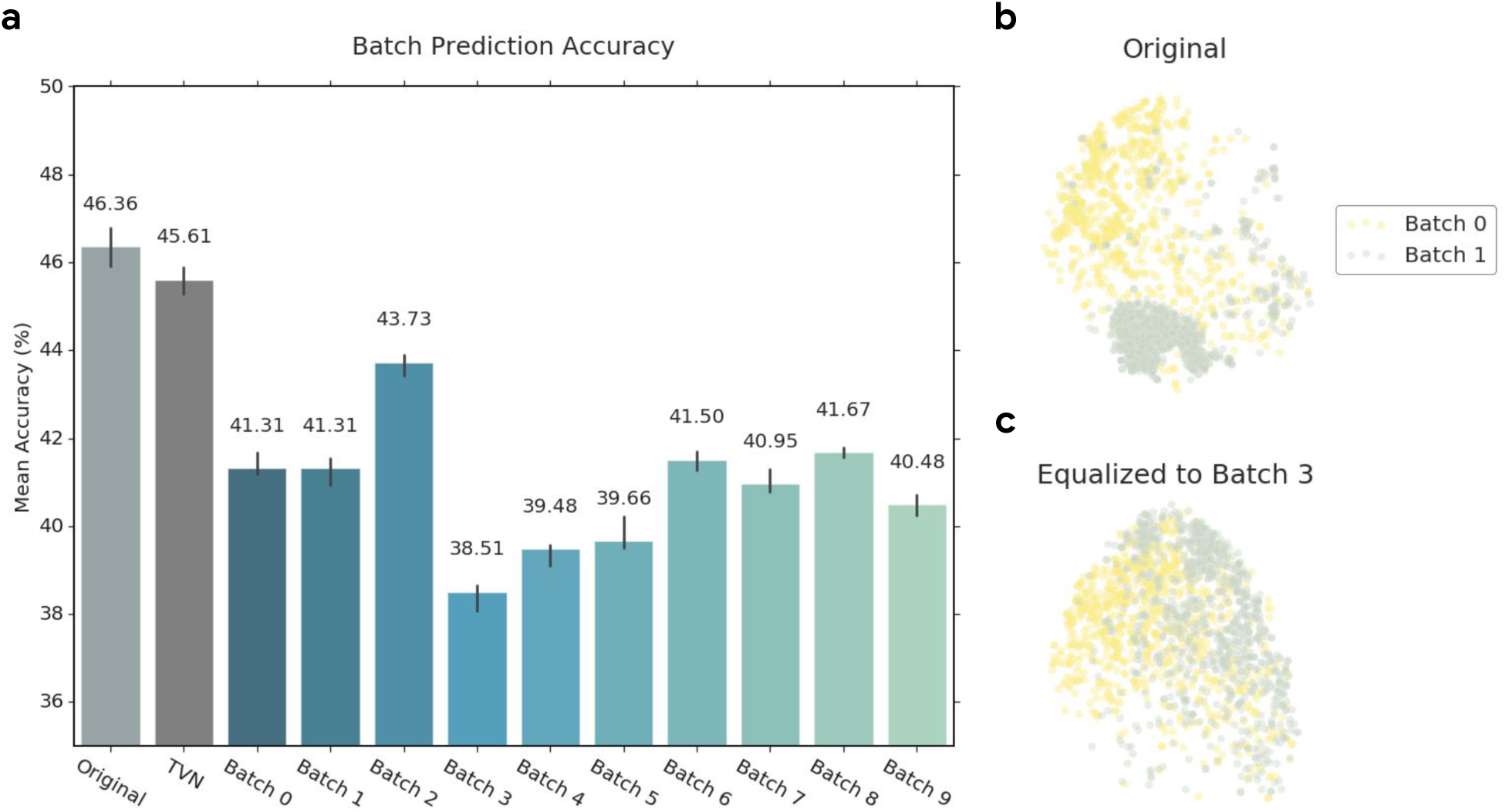
Batch equalization GAN reduces batch variation. a) We equalize the original image dataset to batch 0 through 9 separately and compare their imaging embedding batch prediction accuracy (on the negative control) using a random forest classifier. Error bars indicate the min and max accuracy for the five-fold cross-validation study. b) 2-D visualization of the original image embedding (negative control only) through t-SNE where samples image embedding from batch 0 and batch 1 are shown. c) Same 2-D visualization as b) but using the image embedding from an equalized dataset is shown. We use the dataset where all the images are equalized batch 3 because it has the least batch predictive power, as shown in a).

In addition, after a t-SNE dimension reduction, we show that the equalized image embedding (Figure 4c) are less clustered by batch compared to the original image embedding (Figure 4b). By de-clustering the embedding, downstream analyses relying on the embedding distance measure are more robust to the batch effect. All pair-wise embedding visualizations are in Figure S3 and a quantitative measure of the embedding distance difference is shown in Figure S4.

### 4.4 Batch equalization GAN transforms the image to the target batch correctly

To show our generator *G* is working as intended and indeed equalizing the image to targeted batch, we train a random forest classifier on the original image embedding for batch prediction and compare the confusion matrix when applying the same classifier to the test set of original image embedding vs. equalized image embedding. As shown in Figure 5, after equalizing the dataset to batch 3, the original batch classifier is much more likely to predict an image belong to batch 3 and less likely to predict the original batch as shown in the diagonal line. Figure S5 shows the differences in the confusion matrix when equalizing to other target batches.

**Figure 5:**
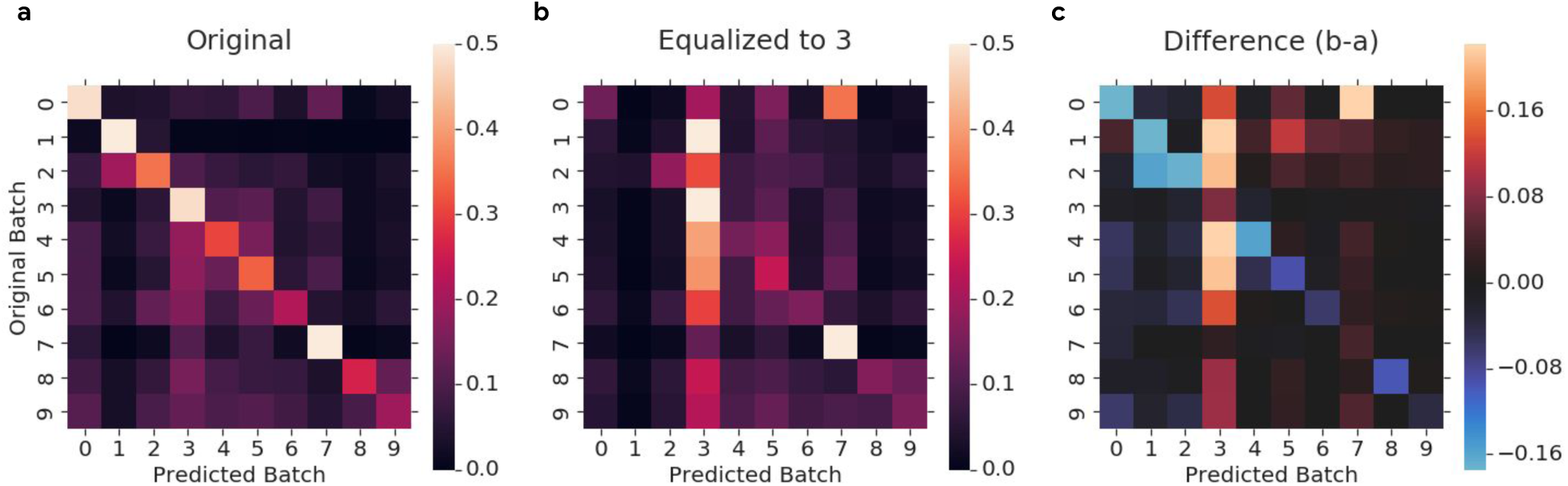
Confusion matrix for batch prediction on the original image embedding. a) The classifier can predict the original batch relatively well with a high probability along the diagonal line. b) Confusion matrix for batch prediction on the equalized to batch 3 image embedding using the same classifier. c) To better highlight the difference after we equalized the image, we show the difference between the confusion matrix after and before we equalized the images to batch 3. The classifier is much more likely to predict an image belong to batch 3, and most of the diagonal line values decrease and show a drop in prediction accuracy.

We also want to highlight that the confusion for batch 7 is not affected by design. Batch 7 has especially high confluency comparing to other batches, but the generator is not able to remove cells from the image after imposing the entangled loss, so we expect the image in batch 7 to be easily classified as batch 7 even after the equalization.

### 4.5 Batch equalization GAN preserves the important biological features

To quantitatively measure the potential modification to the biological features of the image, we use the image embedding generated from the equalized images to predict the compound mechanism of action (MoA). Since MoA is usually inferred from the biological features of the image, our accuracy in MoA prediction could reflect the quality of the biological features in our equalized images. In this evaluation, we compute MoA performance along the same line as Ljosa et al. (2013), where the task is to match each compound treatment (a compound at a specific concentration) to the nearest treatment that is not the same compound (NSC).

As shown by the blue bars in Figure 6, the MoA prediction gets better after we equalized the dataset. The improvement indicates batch equalization GAN has a very minimal impact on the biological features, and because we can correct for the batch effect shown in Figure 4b/c, we can predict the MoA even better using 1 nearest neighbor (1-NN). Since the original image dataset has already been processed through simple heuristic like flat-field correction, the better performance through equalization also shows a more sophisticated method do correct the batch effect better.

**Figure 6:**
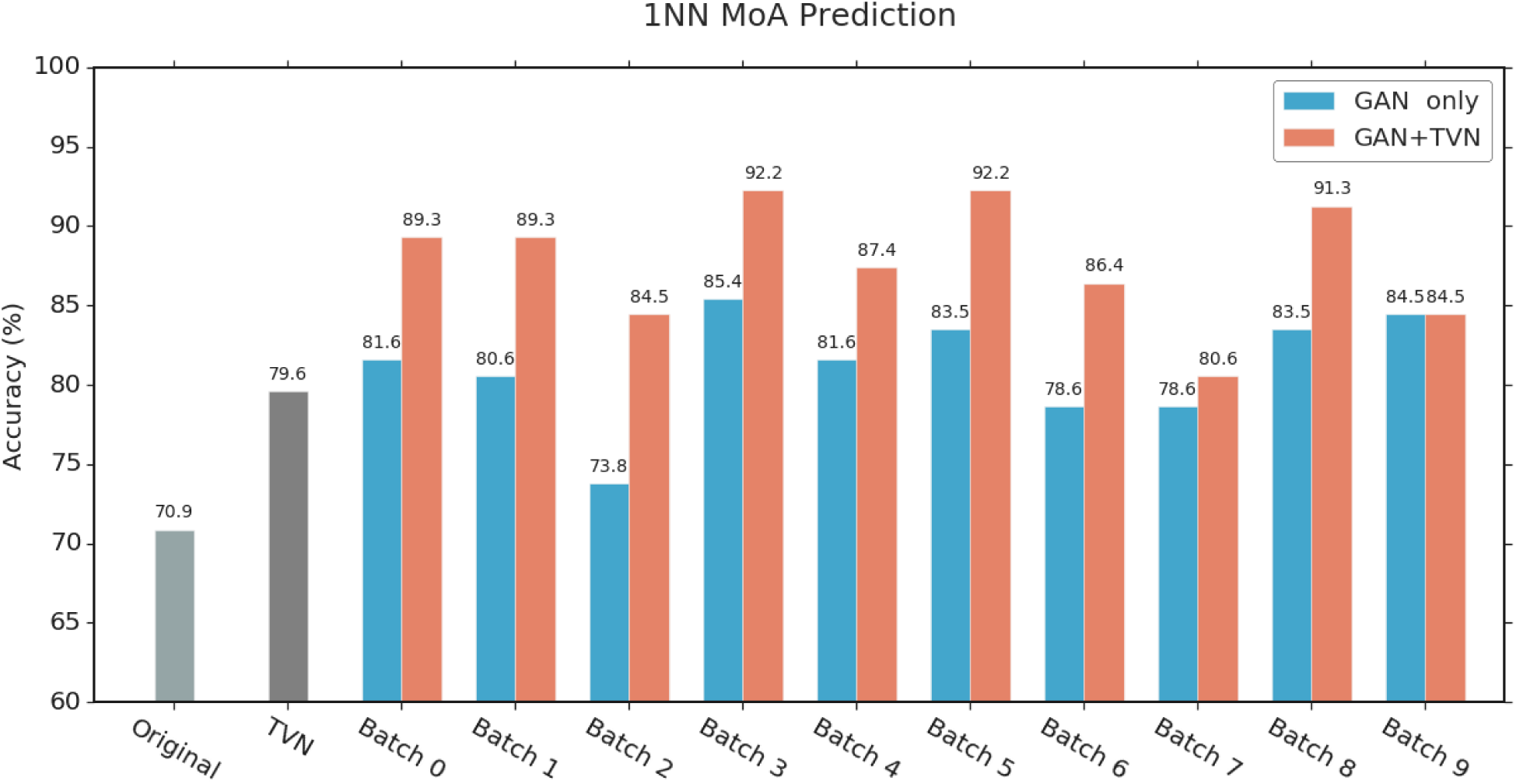
Mechanism of action (MoA) prediction accuracy with 1 nearest neighbor. We equalize the original image dataset to batch 0 through 9 separately and predict the MoA with 1 nearest neighbor using their image embedding. The blue bars represent the MoA prediction accuracy on the equalized image dataset, and the orange bars represent the performance by combining image equalization with TVN embedding normalization.

In addition, we want to point out that the MoA predication accuracy correlates well with the batch prediction accuracy as the equalized dataset that has the least batch predictive power also performs the best here (Figure S6). The correlation shows not only the necessity for batch effect mediation but also that, in the application, we can choose which equalized image dataset or target batch to use for downstream analysis based on their batch predictive power.

### 4.6 Batch equalization GAN can work with other batch normalization method

Since batch equalization GAN can be treated as an image-level pre-processing to mediate the batch effect, it does not affect many of the embedding-level batch normalization methods we used today. In Figure 6, we see similar performance in MoA prediction after applying image equalization or TVN embedding normalization (Ando et al., 2017). However, after combining the two methods (orange bars), we see an even better performance in the MoA prediction comparing to the result when applying each method separately. It indicates that batch equalization in an image level can work well with embedding level normalization methods by mediating different aspects of the batch effect resulting in a stronger performance.

## 5 Discussion

In this work, we introduce batch equalization GAN that allows us to equalize the batch variation by directly modifying the appearance of scientific images. We show that a naive GAN implementation has too much freedom in modifying an image and thus does not preserve the biological semantics of the image. We address this by incorporating an entangled loss extension for the GAN training that preserves the biological semantics while still producing images whose appearance matches the targeted batch. We believe our approach enables a novel analysis for multi-week experiments through the creation of a hypothetical version of the ideal experiment where every experimental condition is tested every week.

However, our work has also revealed limitations in our ability to equalize all experimental variation and extract only the relevant biological information. For instance, in Fig. 5b, our method fails to transform images in batch 7 to batch 3 sufficiently to fool our batch classifier. We also note that our MoA prediction is only able to achieve 92% 1-NN accuracy as opposed to the 97% 1-NN accuracy by Caicedo et al. (2018) that also analyzes the RGB images. We believe that our failure to equalize batch 7 to batch 3 is due to the abnormally high cell density of batch 7. Our batch equalization GAN is trained to *not* change the biological semantics of an image like adding or subtracting cells so that the batch classifier can take advantage of the different cell density for its classification. We also have not optimized our embeddings for the scientific RGB images (e.g., performing semi-supervised training like Caicedo et al. (2018)), so we do not expect to achieve their state-of-the-art performance. Instead, our approach is compatible with any downstream analysis method, so we believe our approach should be able to match existing performance and improve the robustness of comparisons across batches.

The tension between what variation is truly biological rather than technical highlights the power of equalizing the images rather than an abstract representation (e.g., an embedding or another large-dimensional vector of measurements). If the cells are in a different state (e.g., stressed and unhealthy in one week), then they could have different biological responses to perturbation. We want our analysis to capture when experimental conditions are incomparable rather than inappropriately normalizing them to the same value, and the ability to interpret our model’s changes to the data is essential for that building trust.

The ability to generate hypothetical datasets opens up a new possibility for how to use deep learning to understand a dataset. Standard applications of deep learning to microscope image preprocessing are focused on exposing unseen details such as super-resolution (Wang et al., 2019) or image reconstruction (Belthangady and Royer, 2019). However, deep learning may not only be useful in analysis, but it could also be useful in data exploration based on transformations between conditions. Even the naive GAN could be helpful in data exploration by exposing the differences, biological and technical, between conditions that are intended to be the same. These types of ‘what if’ questions are difficult to ask in a large-scale dataset, and we hope these exploratory uses of machine learning enable a new workflow between scientists and algorithms.

## Acknowledgements

We thank Luke Metz for his helpful comments, Joel Shor for his code reviews, and the Google Accelerated Science Team for their helpful discussions and suggestions.

## Supplementary Materials and Methods

### Details for image preprocessing and normalization

For image prepossessing and normalization, we performal similar procedure for bit conversion as Ando et al. (2017) and image normalization as Singh et al. (2014). We first identified cell candidate regions using just the DAPI channel. For all comparisons to previous results, we used with the DAPI center locations of Ljosa et al. (2013) and filtered out any that are too close to the border. We then converted all of the 16-bit integer images into 32-bit float images, so we can perform our preprocessing without introducing errors from integer rounding. We computed a flat field image for each plate and channel similar to Singh et al. (2014), except we use the 10th percentile rather than the median, and we blurred the resulting image with a gaussian sigma of 50. Next, we divided each image by its appropriate flatfield image, producing an image that represents the signal/background at each pixel location. To improve the dynamic range before converting to an 8-bit integer image, we set the minimum value to 1.0 and take the natural log of each pixel value. We then found the minimum and maximum values of the image, and we clipped the maximum to be no greater than 5 to prevent extreme values. We then linearly re-scaled the minimum and maximum to be the values 0 and 255. To create the final 8-bit RGB image, we rounded the floating-point values and convert the values to 8-bit unsigned integers.

### Parameterization for the generator and discriminator

For the parameterization of our generator and discriminator, we use a very similar architecture as the ones described in StarGAN (Choi et al., 2017). For our generator *G*, we use the same architecture as their *G*_StarGAN_. However, since we do not expect a dramatic shift for the edges in cell images, we only use the neural network to generate the modification Δ as *G*(**X**) = **X** + *G*_StarGAN_(**X**) where *G*_StarGAN_(**X**) is only modeling a “painting” on top of the input image. For our discriminator *D*, we use a very similar architecture as their *D*_StarGAN_: 1) instead of six convolutions layers with depths (64, 128, 256, 512, 1024, 2048), we use six convolution layers with 1*/*4 of the depths (16, 32, 64, 128, 256, 512) to speed up the training since the simpler discriminator structure does not affect the overall training; 2) as a new disentanglement term is introduced, we have an additional output layer whose final output is (1, 1, *n*_**Y**_) where *n*_**Y**_ is the number of biological features.

### Hyper-parameters and implementation details

During the training phase, we sample 2 images from the negative control of each of the 10 batches and randomly choose their corresponding targets forming the training data for one iteration (i.e., a batch size of 20). We then train the generator and discriminator function with respect to the objectives *L*_*G*_ and *L*_*D*_ via Adam (*β*_1_ = 0.5 and *β*_2_ = 0.999). We started with a learning rate of 0.0001 for both the generator and discriminator network and let the learning rate decay to 0 linearly after running over 500000 iterations. To stabilize the training, we follow the training procedure in the improved Wasserstein-GAN, where for each iteration, we train the generator for 1 step and the discriminator for 5 steps. For the the objective function, we set the coefficients as follow: *λ*_gp_ = 10, *λ*_batch_ = 1, *λ*_rec_ = 10, and finally *λ*_entangled_ = 25. Our training hyper-parameters closely follow those in StarGAN (Choi et al., 2017), while *λ*_entangled_ is selected from a set of numbers {1, 5, 25, 100} because, qualitatively, it shows the best trade-off between batch transformation and biological feature preservation. The training takes about a day with four NVIDIA Tesla V100 GPUs. We reimplemented the model in Tensorflow, and this general-purpose StarGAN reimplementation is open-sourced by us during the development of this work: https://github.com/tensorflow/gan.

### Image embedding

For our analysis related to the image embedding, we generated a 64-dimensional embedding from the input RGB image using a Deep Metric Network (Wang et al., 2014) pre-trained on consumer image datasets. Since the mapping of the scientific stain channel to the RGB channel is arbitrary, we performed an embedding on all six possible mappings of DAPI, Tubulin, and Actin to the red, green, and blue channels. We then averaged them to form the final embedding of the images.

While our embedding process closely follows Ando et al. (2017), a key difference is that we are merging the three images (DAPI, Tubulin, and Actin) into a single RGB image rather than feeding each as a separate grayscale image through the model. As a consequence, we have a single embedding for an image rather than concatenating three embeddings together. Besides, we used an alternative neural embedding model that is more sensitive to batch variation caused by color after pre-training on consumer images. These embedding with more batch variation gives us more room to demonstrate the reduction of batch effect.

### Random Forest classifier for batch prediction

A random forest classifier is used to predict the batch of an image from its image embedding. We used the default random forest classifier implementation from Scikit-learn (Pedregosa et al., 2011) with the default hyper-parameter.

### t-SNE dimension reduction

To visualize the image embedding in 2-D, we use t-SNE (Maaten and Hinton, 2008) to reduce the dimension of our 64-D image embedding to 2-D. We used the implementation from Scikit-learn (Pedregosa et al., 2011) with all the default hyper-parameter.

**Figure S1:**
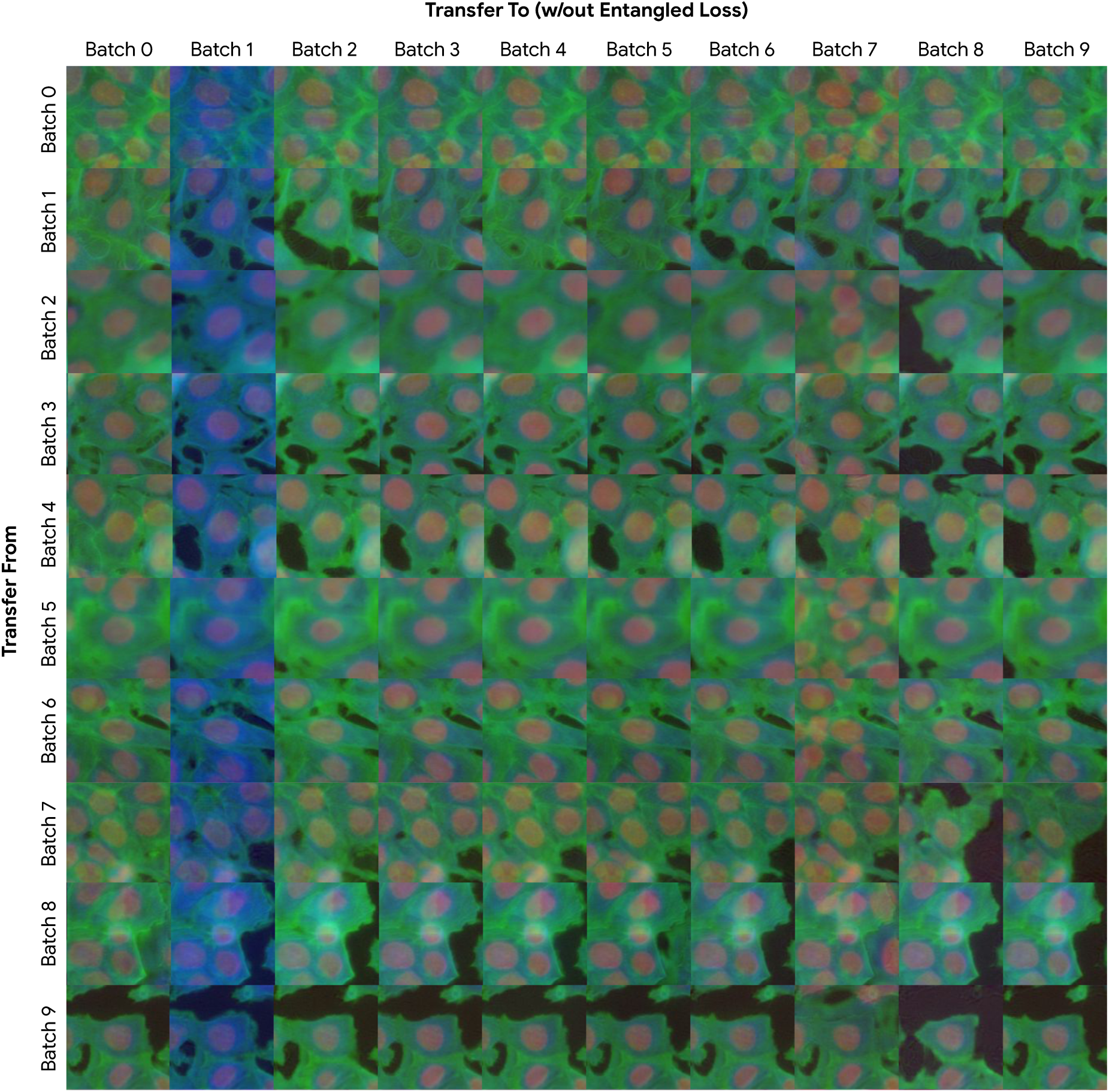
Batch equalization using generator trained without the entangled loss ℒ_entangled_. Corresponding to Fig. 3a. Since we are not restricting the model from changing the biological property, the generator ends up adding cells and reducing the cell size.

**Figure S2:**
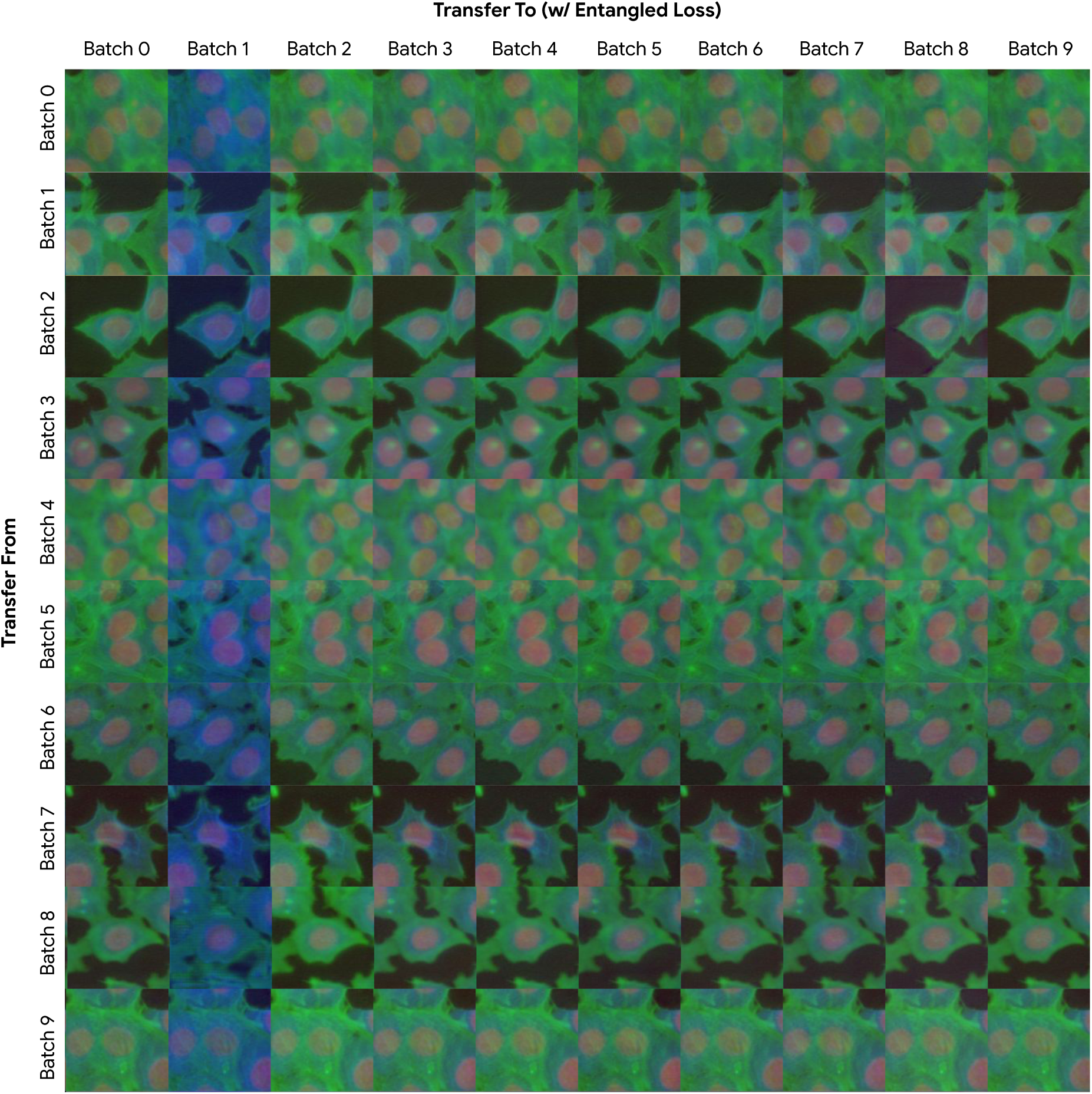
Batch equalization using generator trained with the entangled loss ℒ_entangled_. Corresponding to Fig. 3b. With the entangled loss, the generator is restricted from modifying the biology and only making style changes to the original image.

**Figure S3:**
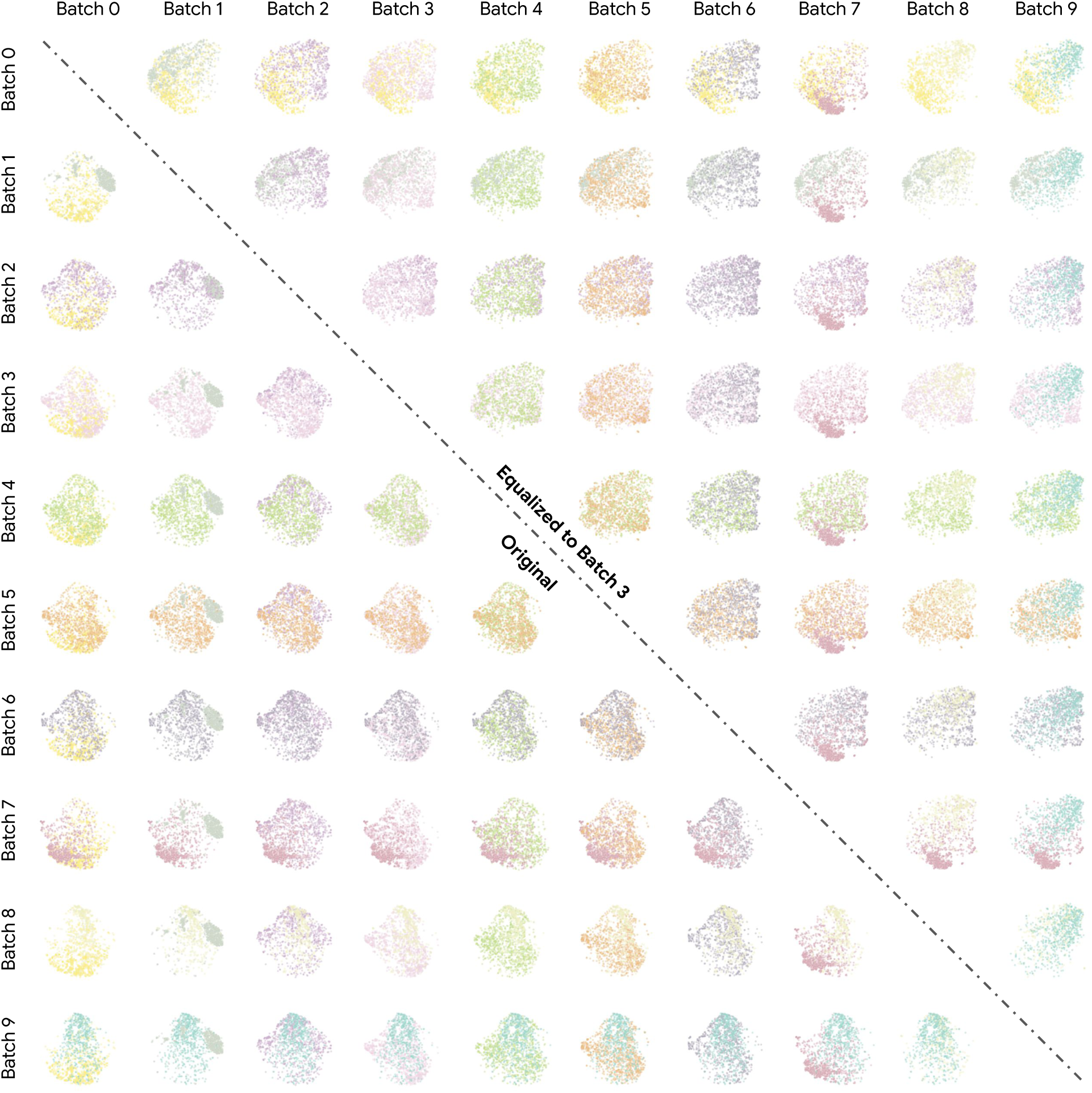
2-D visualization of the image embedding for all pairs of batches. Different colors are representing different batches. The lower left triangle is the original embedding (corresponding to Fig. 4b. The upper right triangle is the embedding after equalizing all images to batch 3 (corresponding to Fig. 4c. The equalization fails to de-cluster batch 7 because our model restricts the generator from making up new cells. For batch 1, whose color distribution is very different, our generator can successfully de-cluster it.

**Figure S4:**
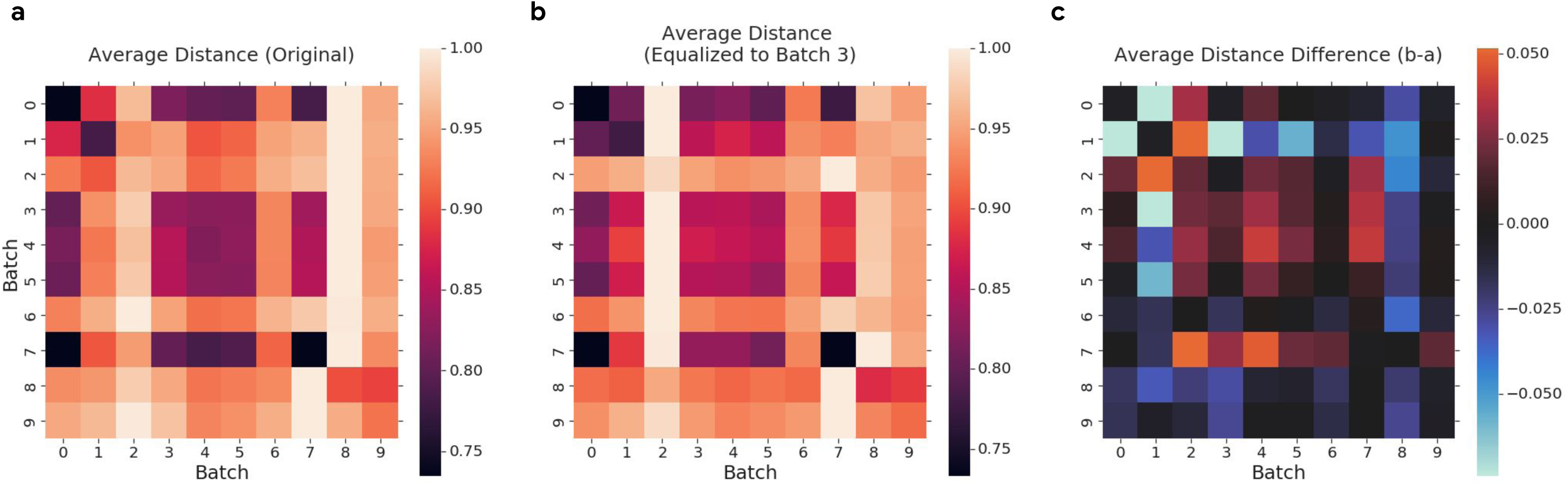
Average batch to batch distance. a) We sampled 500 original image embedding from each batch and calculated their average euclidean distance between each batch. We also normalized the final 10 *×* 10 distance matrix by the largest batch to batch distance so that we have a max distance of 1. b) Same distance matrix as a) but using the image embedding from an equalized dataset is shown. We use the dataset where all the images are equalized batch 3. c) Distance difference between the equalized image embedding and original image embedding. Most of the intra-batch distances get larger, while many inter-batch distances get smaller, indicating a successful de-clustering.

**Figure S5:**
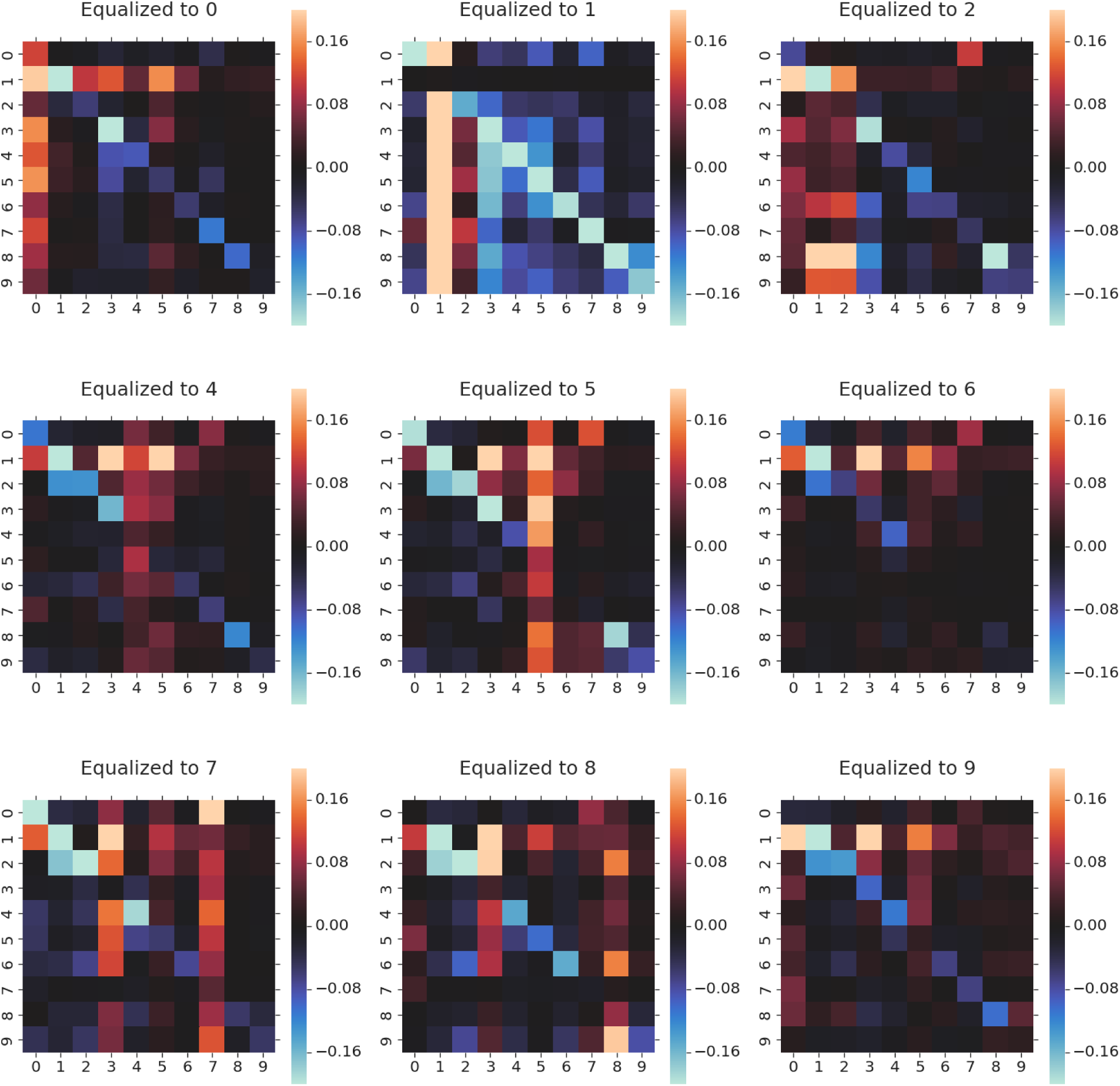
Confusion matrix difference for prediction accuracy. The X-axis is the predicted batch, and the Y-axis is the original batch. This heat map shows how the confusion matrix change after we equalized the image to the corresponding target batch, the same as Fig. 5c. For most of the batch, the generator can fool the classifier to get a high prediction probability for the targeted batch. For the batch that fails to do so (e.g., 6, 9), the prediction accuracy still drops significantly for the original batch.

**Figure S6:**
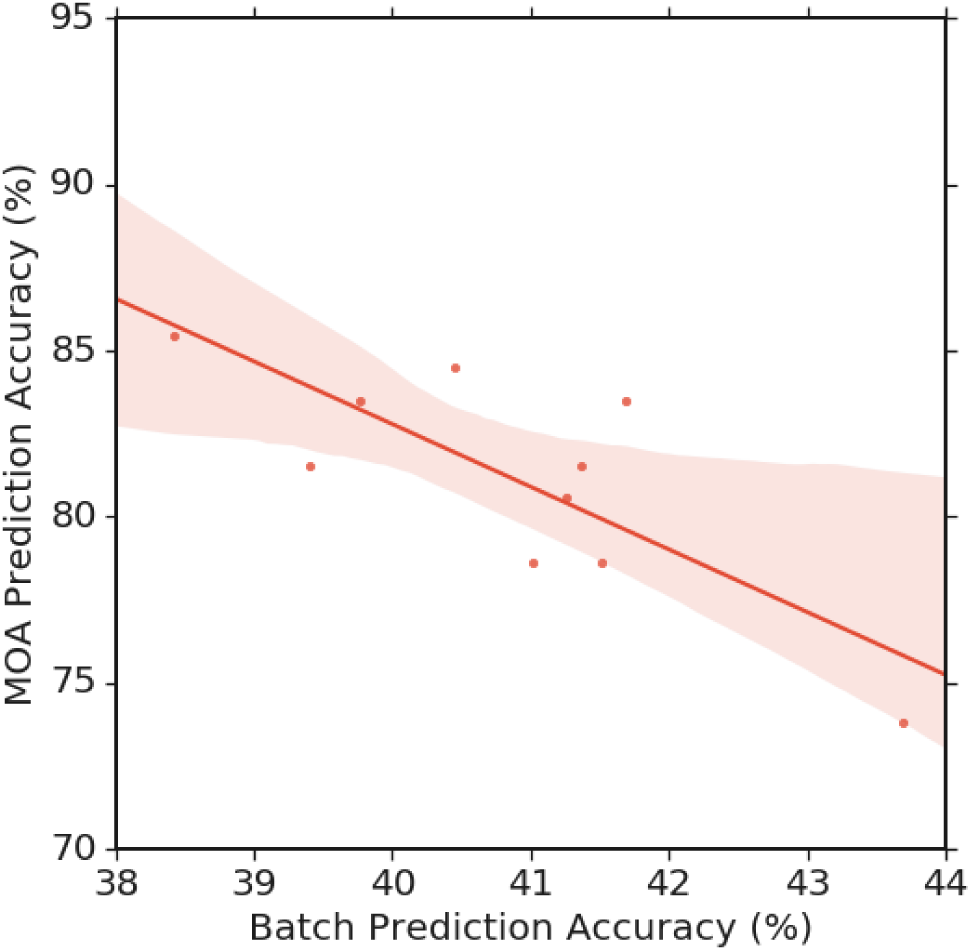
Correlation between the batch prediction accuracy and MOA prediction accuracy. The Pearson correlation coefficient *r* = −0.79.

## References

Amodio, M., van Dijk, D., Montgomery, R., Wolf, G., and Krishnaswamy, S. (2018). Out-of-Sample extrapolation with neuron editing.

Ando, D. M., McLean, C., and Berndl, M. (2017). Improving phenotypic measurements in High-Content imaging screens. bioRxiv, page 161422.

Belthangady, C. and Royer, L. A. (2019). Applications, promises, and pitfalls of deep learning for fluorescence image reconstruction. Nat. Methods.

Caicedo, J. C., Cooper, S., Heigwer, F., Warchal, S., Qiu, P., Molnar, C., Vasilevich, A. S., Barry, J. D., Bansal, H. S., Kraus, O., et al. (2017). Data-analysis strategies for image-based cell profiling. Nature methods, 14(9):849.

Caicedo, J. C., McQuin, C., Goodman, A., Singh, S., and Carpenter, A. E. (2018). Weakly supervised learning of single-cell feature embeddings. In The IEEE Conference on Computer Vision and Pattern Recognition (CVPR).

Caie, P. D., Walls, R. E., Ingleston-Orme, A., Daya, S., Houslay, T., Eagle, R., Roberts, M. E., and Carragher, N. O. (2010). High-content phenotypic profiling of drug response signatures across distinct cancer cells. Mol. Cancer Ther., 9(6):1913–1926.

Choi, Y., Choi, M., Kim, M., Ha, J.-W., Kim, S., and Choo, J. (2017). StarGAN: Unified Generative Adversarial Networks for Multi-Domain Image-to-Image Translation. arXiv e-prints, page 1711.09020.

Goodfellow, I., Pouget-Abadie, J., Mirza, M., Xu, B., Warde-Farley, D., Ozair, S., Courville, A., and Bengio, Y. (2014). Generative adversarial nets. In Ghahramani, Z., Welling, M., Cortes, C., Lawrence, N. D., and Weinberger, K. Q., editors, Advances in Neural Information Processing Systems 27, pages 2672–2680. Curran Associates, Inc.

Gulrajani, I., Ahmed, F., Arjovsky, M., Dumoulin, V., and Courville, A. C. (2017). Improved training of wasserstein gans. In Advances in neural information processing systems, pages 5767–5777.

Isola, P., Zhu, J.-Y., Zhou, T., and Efros, A. A. (2017). Image-to-image translation with conditional adversarial networks. In Proceedings of the IEEE conference on computer vision and pattern recognition, pages 1125–1134.

Kim, T., Cha, M., Kim, H., Lee, J. K., and Kim, J. (2017). Learning to discover Cross-Domain relations with generative adversarial networks.

Kingma, D. P. and Ba, J. (2014). Adam: A method for stochastic optimization. arXiv preprint 1412.6980.

Kothari, S., Phan, J. H., Moffitt, R. A., Stokes, T. H., Hassberger, S. E., Chaudry, Q., Young, A. N., and Wang, M. D. (2011). Automatic batch-invariant color segmentation of histological cancer images. Proc. IEEE Int. Symp. Biomed. Imaging, 2011:657–660.

Leek, J. T., Scharpf, R. B., Bravo, H. C., Simcha, D., Langmead, B., Johnson, W. E., Geman, D., Baggerly, K., and Irizarry, R. A. (2010). Tackling the widespread and critical impact of batch effects in high-throughput data. Nature Reviews Genetics, 11(10):733.

Liu, A. H., Liu, Y.-C., Yeh, Y.-Y., and Wang, Y.-C. F. (2018). A unified feature disentangler for multi-domain image translation and manipulation. In Advances in neural information processing systems, pages 2590–2599.

Ljosa, V., Caie, P. D., Ter Horst, R., Sokolnicki, K. L., Jenkins, E. L., Daya, S., Roberts, M. E., Jones, T. R., Singh, S., Genovesio, A., Clemons, P. A., Carragher, N. O., and Carpenter, A. E. (2013). Comparison of methods for image-based profiling of cellular morphological responses to small-molecule treatment. J. Biomol. Screen., 18(10):1321–1329.

Maaten, L. v. d. and Hinton, G. (2008). Visualizing data using t-sne. Journal of machine learning research, 9(Nov):2579–2605.

Mirza, M. and Osindero, S. (2014). Conditional generative adversarial nets.

Otsu, N. (1979). A threshold selection method from Gray-Level histograms. IEEE Trans. Syst. Man Cybern., 9(1):62–66.

Pawlowski, N., Caicedo, J. C., Singh, S., Carpenter, A. E., and Storkey, A. (2016). Automating morphological profiling with generic deep convolutional networks. bioRxiv, page 085118.

Pedregosa, F., Varoquaux, G., Gramfort, A., Michel, V., Thirion, B., Grisel, O., Blondel, M., Prettenhofer, P., Weiss, R., Dubourg, V., Vanderplas, J., Passos, A., Cournapeau, D., Brucher, M., Perrot, M., and Duchesnay, E. (2011). Scikit-learn: Machine learning in Python. Journal of Machine Learning Research, 12:2825–2830.

Singh, S., Bray, M.-A., Jones, T. R., and Carpenter, A. E. (2014). Pipeline for illumination correction of images for high-throughput microscopy. J. Microsc., 256(3):231–236.

Tabak, G., Fan, M., Yang, S. J., Hoyer, S., and Davis, G. (2017). Correcting nuisance variation using wasserstein distance.

Venugopalan, S., Narayanaswamy, A., Yang, S., Gerashcenko, A., Lipnick, S., Makhortova, N., Hawrot, J., Marques, C., Pereira, J., Brenner, M., et al. (2019). It’s easy to fool yourself: Case studies on identifying bias and confounding in bio-medical datasets. arXiv preprint 1912.07661.

Wang, H., Rivenson, Y., Jin, Y., Wei, Z., Gao, R., Günaydin, H., Bentolila, L. A., Kural, C., and Ozcan, A. (2019). Deep learning enables cross-modality super-resolution in fluorescence microscopy. Nat. Methods, 16(1):103–110.

Wang, J., Song, Y., Leung, T., Rosenberg, C., Wang, J., Philbin, J., Chen, B., and Wu, Y. (2014). Learning fine-grained image similarity with deep ranking. In Proceedings of the IEEE Conference on Computer Vision and Pattern Recognition, pages 1386–1393.

Zhu, J.-Y., Park, T., Isola, P., and Efros, A. A. (2017). Unpaired Image-to-Image translation using Cycle-Consistent adversarial networks.

